# Dynamic Architecture of DNA Repair Complexes and the Synaptonemal Complex at Sites of Meiotic Recombination

**DOI:** 10.1101/206953

**Authors:** Alexander Woglar, Anne M. Villeneuve

## Abstract

Meiotic double-strand breaks (DSBs) are generated and repaired in a highly regulated manner to ensure formation of crossovers (COs) while also enabling efficient non-CO repair to restore genome integrity. Here we use Structured-Illumination Microscopy to investigate the dynamic architecture of DSB repair complexes at meiotic recombination sites in relationship to the synaptonemal complex (SC). DSBs resected at both ends are rapidly converted into inter-homolog repair intermediates harboring two populations of BLM helicase and RPA, flanking a single population of MutS *γ*. These intermediates accumulate until late pachytene, when repair proteins disappear from non-CO sites and CO-designated sites become enveloped by SC-central region proteins, acquire a second MutS *γ* population, and lose RPA. These and other data suggest that the SC protects CO intermediates from being dismantled inappropriately and promotes step-wise CO maturation by generating a transient CO-specific repair compartment, thereby enabling differential timing and outcome of repair at CO and non-CO sites

## Introduction

During meiosis in most organisms, each pair of homologous chromosomes must receive at least one crossover (CO) in order to undergo reliable segregation during the first meiotic division, thereby preserving euploidy in the next generation. COs are generated by a specialized form of recombinational repair of double–strand DNA breaks (DSBs) introduced by the meiosis-specific SPO-11 protein. In order to ensure that every homolog pair receives at least one CO, a surplus of DSBs is introduced during meiotic prophase. Only a subset of these DSBs mature into COs, while the rest are repaired by alternative homologous recombination-based mechanisms that do not yield COs. Despite an excess of DSBs, COs exhibit wide spacing along the lengths of chromosomes, and many plant and animal models typically undergo only a single CO per chromosome arm. Thus, CO formation is both ensured and robustly limited at the same time (For reviews see: (Mercier et al., 2015; Zickler and Kleckner, 2015, 2016)).

Several conserved meiosis-specific proteins, such as the MutSy (MSH4-MSH5) complex or the cyclin-related protein COSA-1/CNTD1, were identified as being essential for controlled CO formation. These pro-CO factors localize to sites of ongoing meiotic recombination and modulate the outcome of DNA repair to yield COs as final DSB repair products at selected “CO-designated” sites (for review see: (Hillers et al., 2017; Zickler and Kleckner, 2015, 2016). Nevertheless, how these proteins promote the CO repair outcome, and how different modes of DSB repair can operate at other sites on the same chromosomes in the same cells to yield non-CO products, are far from being understood.

Controlled CO formation further depends on components of the synaptonemal complex central region (SC-CR). The SC-CR is composed of predicted coiled-coil domain proteins and assembles between the axes of aligned homologous chromosomes, maintaining them at a distance of roughly 100-150 nm along their lengths throughout the pachytene stage of meiotic prophase. The SC-CR is dispensable for initial homologous pairing and DSB formation, but when it is absent, meiotic COs are reduced or eliminated (depending on the organism), indicating that SC-CR components play a role in promoting CO formation (for review see: (Hillers et al., 2017; Page and Hawley, 2004)). The SC is a long known cytological hallmark of meiosis (Fawcett, 1956; Moses, 1956) and has been implicated previously in multiple aspects of CO regulation (Hayashi et al., 2010; Libuda et al., 2013; MacQueen et al., 2002; Sym et al., 1993; Sym and Roeder, 1994; Voelkel-Meiman et al., 2016). Further, recent studies have revealed dynamic properties of the SC and have begun to provide insight regarding how changes in SC dynamics might contribute both to nucleus-wide and chromosome-autonomous signaling systems that limit DSBs and inhibit excess COs (Machovina et al., 2016; Nadarajan et al., 2017; Pattabiraman et al., 2017; Rog et al., 2017; Subramanian et al., 2016). However, how the SC-CR (or its constituent proteins) functions in promoting CO formation remains poorly understood.

In the nematode *C. elegans*, in contrast to many other model systems, assembly of the SC-CR is not dependent on recombination (Baudat et al., 2000; Dernburg et al., 1998; Giroux et al., 1989; Romanienko and Camerini-Otero, 2000). This makes it possible to differentiate experimentally between the contributions of DNA repair factors and the contributions of SC components to the process of meiotic recombination. In the current work, we exploit this and other features of the *C. elegans* system to investigate how meiotic DNA repair proteins and meiotic chromosome structure collaborate to bring about a robust outcome of meiosis, *i.e.*, formation of a single CO per chromosome pair and restoration of chromosome integrity prior to the meiotic cell divisions (Hillers et al., 2017).

Unlike most previous studies of *C. elegans* meiosis, in this work we employed a detergent-based chromosome spreading protocol (Loidl et al., 1991; Pattabiraman et al., 2017) that affords greatly improved detection of recombination sites. Using this approach, we were able to establish a comprehensive time course of the dynamic localization of meiosis-specific recombination factors and DSB repair proteins over the course of wild-type and mutant meiosis. Further, we used 3D-Structural Illumination microscopy (SIM) to reveal spatial organization of DSB repair complexes at recombination sites and changes in the architecture of these sites during the course of meiotic prophase, allowing us to make inferences regarding the timing of recombination progression and the nature of the underlying DNA intermediates. Moreover, we discovered a unique spatial arrangement of SC-CR components surrounding CO-designated recombination sites that provides a new framework for thinking about how distinct outcomes of DSB repair can be accomplished at different sites within a single cell.

## Results

### Progressive differentiation of meiotic recombination sites during *C. elegans* meiosis

We re-examined the dynamic localization of multiple DSB repair proteins at the sites of ongoing recombination during the course of meiotic prophase in *C. elegans*, using a nuclear spreading protocol that enables improved visualization of these events (Loidl et al., 1991; Pattabiraman et al., 2017). Whereas meiotic chromosome axis components and chromosome-bound recombination/repair proteins are retained in these preparations, most of the nucleoplasmic protein pools that obscure recombination foci are washed out, resulting in superior detection relative to previously-used *in situ* methods (Fig S1A). At the same time, the relative spatial organization of nuclei within a gonad is maintained largely intact, preserving the spatial/temporal gradient of nuclei progressing through meiotic prophase that is a major strength of this experimental system.

Fig 1A and Fig S1B show simultaneous visualization of RAD-51, the sole *C. elegans* DNA strand exchange protein, and RPA-1, a subunit of the eukaryotic ssDNA binding protein RPA. Similar to previous reports (Alpi et al., 2003; Colaiacovo et al., 2003), RAD-51 foci are detected beginning in zygotene, rise in abundance and reach a plateau level during early pachytene before declining and then disappearing during late pachytene (identified based on loss of the early prophase marker DSB-2 (Fig S1B; (Rosu et al., 2013)). In contrast to previous reports in which very few RPA foci were detected during wild-type meiosis (Garcia-Muse and Boulton, 2005), our analysis provided a more comprehensive view of association of RPA with recombination intermediates. RPA foci are initially detected in zygotene and rise in abundance in parallel with RAD-51, but RPA foci continue to increase in abundance after RAD-51 foci have peaked and begun to decline, and RPA foci accumulate to much higher levels, plateauing at approximately 30 foci/ nucleus by the end of early pachytene. Further, whereas the majority of RAD-51 foci in early prophase do not colocalize with RPA, after RAD-51 foci begin to decline, the majority do colocalize with RPA. Moreover, whereas RPA foci decline markedly at the early-to-late pachytene transition, RPA remains detectable at a substantial number of recombination sites before eventually being lost from all sites in late pachytene.

**Fig 1.**
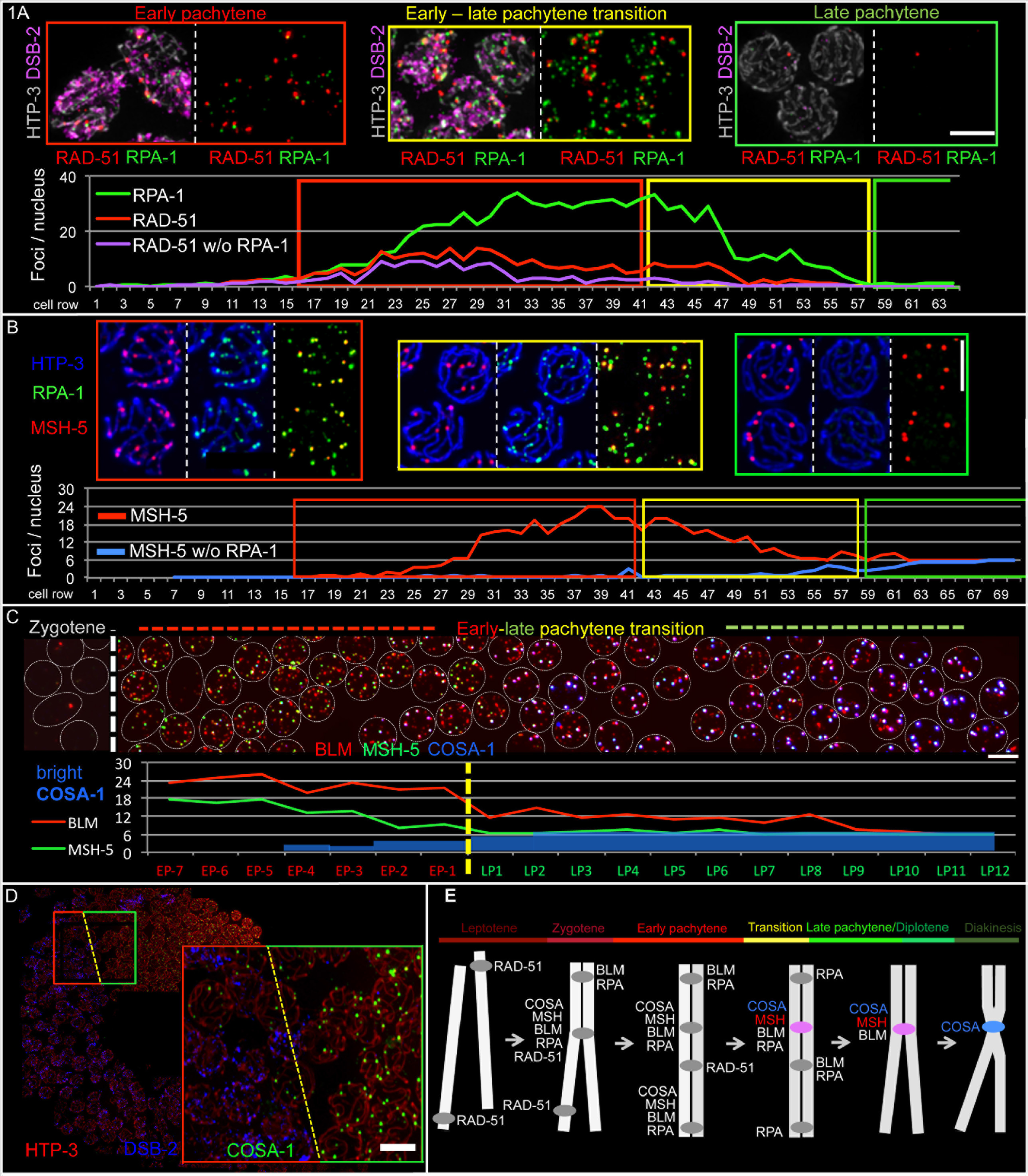
Dynamic localization of RAD-51, RPA, BLM, MSH-5 and COSA-1 over the time course of meiotic prophase (see also Figure S1 and Figure S2) **(A)** Top: Representative enlargements of different meiotic stages of a spread WT gonad (see Fig S1B) probed for RAD-51 (red) and RPA-1 (green). Presence of DSB-2 (purple) indicates nuclei in early prophase stages, and HTP-3 (grey) marks meiotic chromosome axes. Bottom: Quantitative display of appearance, disappearance and co-localization of RAD-51 and RPA-1 foci in a single spread gonad (distinct from the example depicted). **(B)** Top: Representative enlargements of a spread WT gonad probed for RPA-1 (green), MSH-5 (red) and HTP-3 (blue). Bottom: Quantitative display of appearance, disappearance and co-localization of MSH-5 with RPA-1 foci in a single spread gonad (distinct from the example depicted). **(C)** Top: Enlargement of the early-to-late pachytene transition of a spread gonad, probed for GFP::COSA-1 (blue), BLM::HA (red), MSH-5 (green) and HTP-3 (see Fig S2A). Individual nuclei are represented by white circles. Bottom: Quantitative display of appearance and disappearance of BLM, MSH-5 and bright COSA-1 foci at the early-to-late pachytene transition section of the gonad shown above. **(D)** Spread gonad, probed for COSA-1 (green), DSB-2 (blue) and HTP-3 (red). The enlargement on the right highlights the early (red outline) to-late (green outline) pachytene transition, coinciding with the appearance of six bright COSA-1 foci. Scale bars in A-D represent 5 μm. **(E)** Schematic time course of appearance and disappearance of recombination factors at recombination sites during meiotic prophase of *C. elegans*.

Our experiments also revealed substantial overlap between early prophase RPA-marked recombination intermediates and sites of accumulation of MSH-5 (Fig 1B). MSH-5 and MSH-4 make up the heterodimeric MutSγ complex that concentrates at CO-designated sites in late pachytene and is essential for CO formation in *C. elegans* (Hollingsworth et al., 1995; Kelly et al., 2000; Ross-Macdonald and Roeder, 1994; Yokoo et al., 2012; Zalevsky et al., 1999). MSH-5 foci first appear in zygotene, and by the end of early pachytene, nuclei have accumulated 15-25 MSH-5 foci. At this stage, MSH-5 foci nearly always colocalize with RPA foci, with fewer than one RPA-negative MSH-5 focus per nucleus. MSH-5 foci represent interhomolog (IH) recombination intermediates, as they are usually not detected at DNA repair sites on chromosome segments that are unable to engage a (non-sister) homologous chromosome segment as a template for HR (Fig S1C-D). At the early-to-late pachytene transition, MSH-5 is lost from the majority of repair sites while being retained at a single CO-designated repair site on each chromosome pair. These late MSH-5 are brighter than the earlier foci, suggesting an increased local concentration of MutSγ at CO-designated sites. RPA foci also decrease in number following the transition to late pachytene, but in contrast to MSH-5, RPA resides longer at (MSH-5-negative) NCO sites and vanishes from (MSH-5-positive) future CO sites, suggesting a transition at CO sites to a recombination intermediate that lacks ssDNA.

Throughout prophase MSH-5-marked sites also harbor BLM helicase (Fig 1C and Fig S2A). BLM (aka HIM-6 in worms) is required for efficient CO maturation at CO-designated sites, but has also been implicated in dismantling intermediates and completing repair at sites not designated to become COs (for review see: (Hatkevich and Sekelsky, 2017). All BLM foci in early prophase localize with RPA; in contrast, RPA foci lacking BLM are also detected, predominantly at the early-to-late pachytene transition. Similar to RPA, BLM temporarily remains at NCO repair sites after removal of MSH-5. In contrast to RPA, BLM is retained at CO sites throughout late pachytene (similar to MSH-5), while RPA is lost (Fig 1B and Fig S2B).

Previous work had identified the conserved cyclin-related protein COSA-1 as a marker of CO sites in *C. elegans* meiosis (Yokoo et al., 2012). Our spread preparations reveal that in addition to becoming highly concentrated together with MSH-5 at CO sites in late pachytene nuclei, GFP::COSA-1 also decorates a subset of recombination sites in early prophase as very faint foci, co-localizing with MSH-5, BLM and RPA (Fig 1C and S2). These foci are more numerous and appear substantially smaller than the six bright COSA-1 foci detected at CO sites in late pachytene. Loss of COSA-1 and MSH-5 from NCO sites and their accumulation on CO sites in late pachytene occurs concomitantly (Fig 1C). Further, we observe that loss of early prophase nuclear marker DSB-2 strongly coincides with the presence of 6 bright COSA-1 foci, and conversely, that if nuclei retain DSB-2, they invariably have fewer than 6 (if any) bright COSA-1 foci (Fig 1D). This observation indicates an abrupt change in the state of recombination sites as germ cells transition to the late pachytene stage.

Based on these combined observations and previous studies, the following picture emerges (Fig 1E): DSBs are introduced, resected and load RAD-51 throughout early prophase. RAD-51-decorated DNA ends engage in homology search and strand exchange and turn over into RAD-51-free intermediates that contain RPA and BLM. A subset of IH repair sites additionally recruits MutSγ (and COSA-1) to create intermediates with the potential to become COs. Multiple such sites accumulate until each chromosome pair has received at least one such IH intermediated event before nuclei transition to the late pachytene stage. A single DSB repair site on each chromosome pair is designated to become the future CO, either before or during a rapid transition to late pachytene (Rosu et al., 2011; Rosu et al., 2013; Stamper et al., 2013; Woglar et al., 2013). Upon transition to late pachytene, recombination sites designated to mature as COs or NCOs become differentiated from each other, with COSA-1 and MSH-5 becoming enriched at future CO sites, while they are lost from NCO sites at the same time. BLM and RPA remain associated with NCO sites briefly after the transition to late pachytene but disappear soon thereafter, presumably reflecting earlier completion of NCO repair at these sites. Concomitantly, CO-designated sites lose RPA, indicating completion of repair DNA synthesis at those sites; these differentiated CO sites persist throughout the late pachytene stage and vanish by diakinesis, when the presence of a chiasma connecting each homolog pair reflects the presence of a completed CO event.

### The architecture of recombination sites during early prophase

Having established a comprehensive time course of dynamic localization of DNA repair proteins at recombination sites, we combined nuclear spreading with 3D Structured Illumination Microscopy (SIM) to better understand how recombination sites are organized spatially and how this organization changes during meiotic progression. As these spread preparations both greatly improve signal-to-noise ratio and decrease sample thickness, they optimize the specimens for SIM analysis. This enabled us to visualize the relative spatial organization of different recombination proteins present at CO and NCO sites and their relationships to chromosome axes and the SC-CR.

#### RAD-51

SIM imaging revealed that RAD-51 foci usually occur as doublets (Fig 2A). RAD-51 signals are readily resolved as pairs of closely associated foci in the majority of cases in 2D projections of SIM images, and in >70% of cases when SIM images are analyzed in 3D; in the remaining cases, RAD-51 is detected either as an elongated focus or as a single point focus. RAD-51 signals are detected as doublets regardless of the number of DSB sites present in a nucleus and also when chromosomes are strongly spread, and furthermore, we never detect more than two foci at a single site. Also, RAD-51 signals (at least one of the two foci in a pair) are consistently found in close association with chromosome axis markers. Together these data indicate that RAD-51 doublets represent single DSB events (rather than two closely adjacent DSBs) and that both sides of a DSB are resected and load RAD-51. This suggests that both ends are potentially capable of promoting homology search and strand invasion during *C. elegans* meiosis, as was recently shown for budding yeast (Brown et al., 2015).

**Fig 2.**
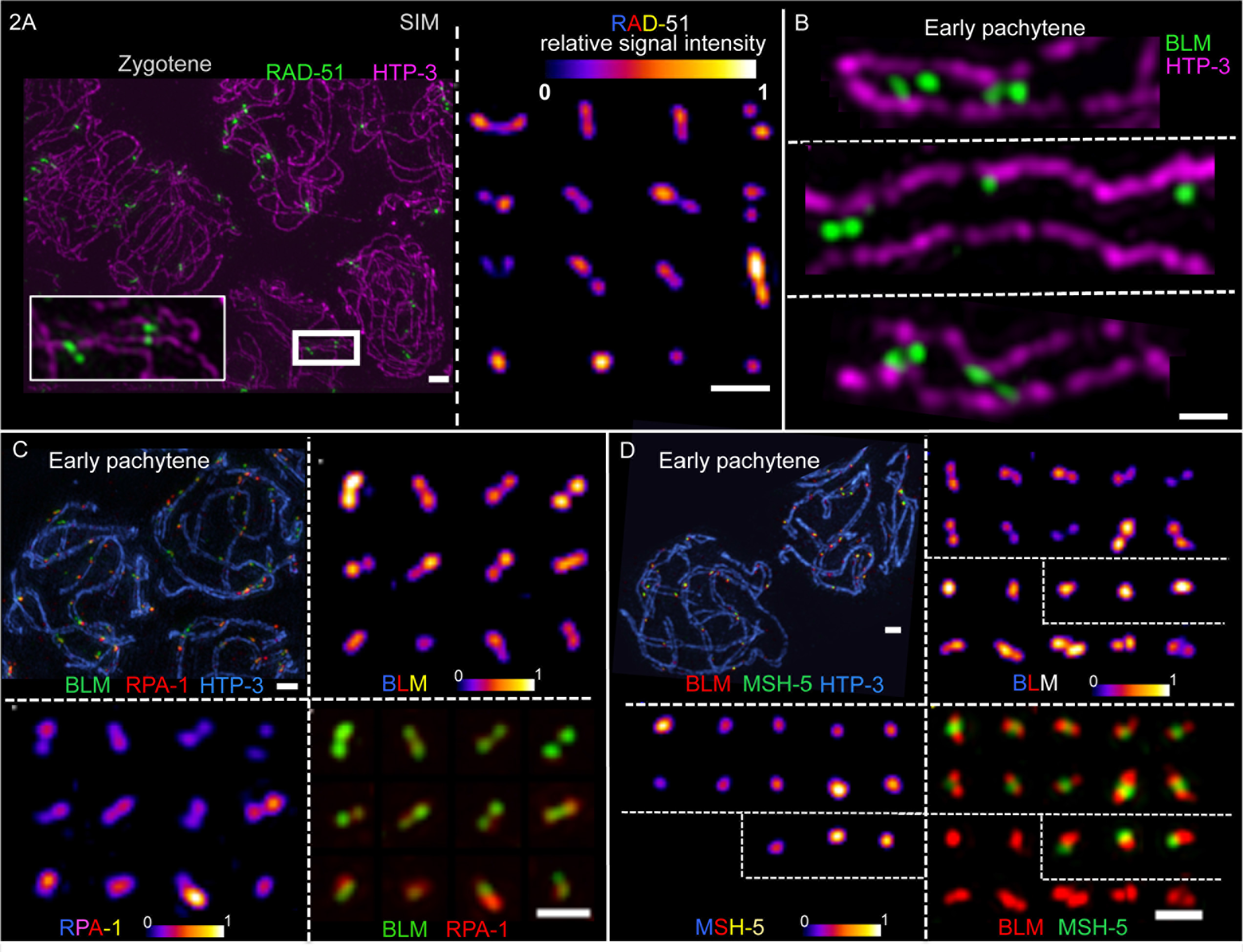
Structural features of DNA repair complexes at recombination sites during early meiotic prophase. Images of nuclei, chromosome segments, and individual recombination sites generated using 3D Structured-Illumination Microscopy (SIM) and Z-stack projection. **(A)** Left panel: WT zygotene nuclei probed for HTP-3 (purple) and RAD-51 (green). The enlargement on the bottom highlights that doublets of RAD-51 are found in close proximity to, but without special orientation relative to chromosome axes. The scale bar represents 1 μm. Right panel: Examples of individually cropped RAD-51 signals (from the same SIM field, as the example on the left). To illustrate the structural organization of individual foci, relative signal intensities are displayed using an 8-bit color look-up table (“LUT Fire” in ImageJ), representing lower intensities in “colder” colors and higher signal intensities in “warmer” colors as indicated. Scale bar: 500 nm. **(B)** Early pachytene nuclei, probed for BLM::HA (green) and HTP-3 (purple). BLM is resolvable either as single foci or as doublet focus throughout prophase. For some segments of a synapsed bivalent where the SCs are oriented in a full frontal view, the two homolog axes are resolvable as two parallel tracks in XY; three such examples are individually cropped and displayed to illustrate the roughly parallel orientation of BLM doublet foci between the parallel-aligned axes. Scale bar: 500 nm. **(C)** Early pachytene nuclei, probed for RPA-1 (red), BLM::HA (green) and HTP-3 (blue). RPA-1 single and double foci are found both with and without BLM. When RPA-1 is resolvable as a doublet, the two foci are co-oriented with the pair of BLM foci, but are slightly off-set. The individually cropped foci originate from the same SIM field as the representative nuclei displayed on the top left. **(D)** Early pachytene nuclei, probed for BLM (red), MSH-5 (green) and HTP-3 (blue). BLM single and double foci are found both with, and without, a single MSH-5 focus in their middle during early prophase. MSH-5 is not resolvable as two foci during early prophase and decorates a different domain than BLM at the same DNA repair site. The individually cropped RPA-1 and BLM foci originate from the same SIM field as the two nuclei in the top left panel. Scale bars in C and D: 1 μm (top left) and 500 nm (bottom right).

#### BLM

We recently reported that BLM is frequently detected as dual foci aligned in parallel with the chromosome axis in pachytene nuclei analyzed *in situ* in whole-mount preparations of *C. elegans* gonads (Jagut et al., 2016). We confirm and extend these findings here for BLM using SIM on nuclear spreads (Fig 2B). In well-spread early prophase nuclei, we can clearly resolve two populations of BLM at up to 75% of recombination sites; these two BLM populations are arranged longitudinally along the chromosomes, and when the axes can be resolved as two parallel tracks, BLM signals are located between the two axial tracks. Furthermore, multiple BLM double and single foci can be observed between the axes along the length of a single pair of homologous chromosomes (bivalent) during early pachytene.

#### RPA

As BLM consistently co-localizes with RPA during zygotene and early pachytene, we investigated the spatial relationship between BLM and RPA signals at the sites of these presumed post-strand invasion repair intermediates. Similarly to BLM, RPA is also typically detected as two foci oriented longitudinally along the chromosomes and between the axes (Fig 2C). However, BLM and RPA don’t strictly colocalize, as RPA sometimes localizes as a single focus in the middle of a BLM doublet. Further, when both RPA and BLM are detected as doublets, their relative orientation is similar, yet slightly oblique. This suggests that ssDNA (marked by RPA) and BLM helicase complexes occupy nearby but distinct positions within the recombination intermediates.

#### MSH-5

At IH recombination sites marked by MSH-5 during zygotene and early pachytene, MSH-5 is consistently detected as a single focus, located either in the middle of a BLM/RPA doublet or side-by-side with a BLM/RPA singlet. Detection of BLM or RPA as doublets does not depend on the presence of MSH-5 at a recombination site, as BLM or RPA doublets are readily detected both at MSH-5-positive and MSH-5-negative recombination sites (Fig 2D).

### The architecture of CO sites during late prophase

SIM images of hypotonic spreads revealed that CO-designated recombination sites acquire a striking new architecture following transition to late pachytene. CO sites not only lose RPA (presumably reflecting completion of repair synthesis), they also change the spatial organization of proteins that remain and/or become concentrated at these sites. As in early pachytene, BLM is still present at each CO repair site as two foci aligned in parallel with and between the homolog axes (Fig 3A). However, in contrast to the MSH-5 singlet signals present in early pachytene, MSH-5 signals at CO sites are now reliably resolved as two foci, oriented perpendicular to and spanning the distance between the aligned homolog axes (Fig 3A). The COSA-1 signal, which increases in intensity over the course of late pachytene, is detected at the center of the cruciform structure defined by the orthogonally-oriented MSH-5 and BLM doublets (Fig 3A). We infer that this organization reflects the presence of a late CO-specific recombination intermediate at these sites.

**Fig 3.**
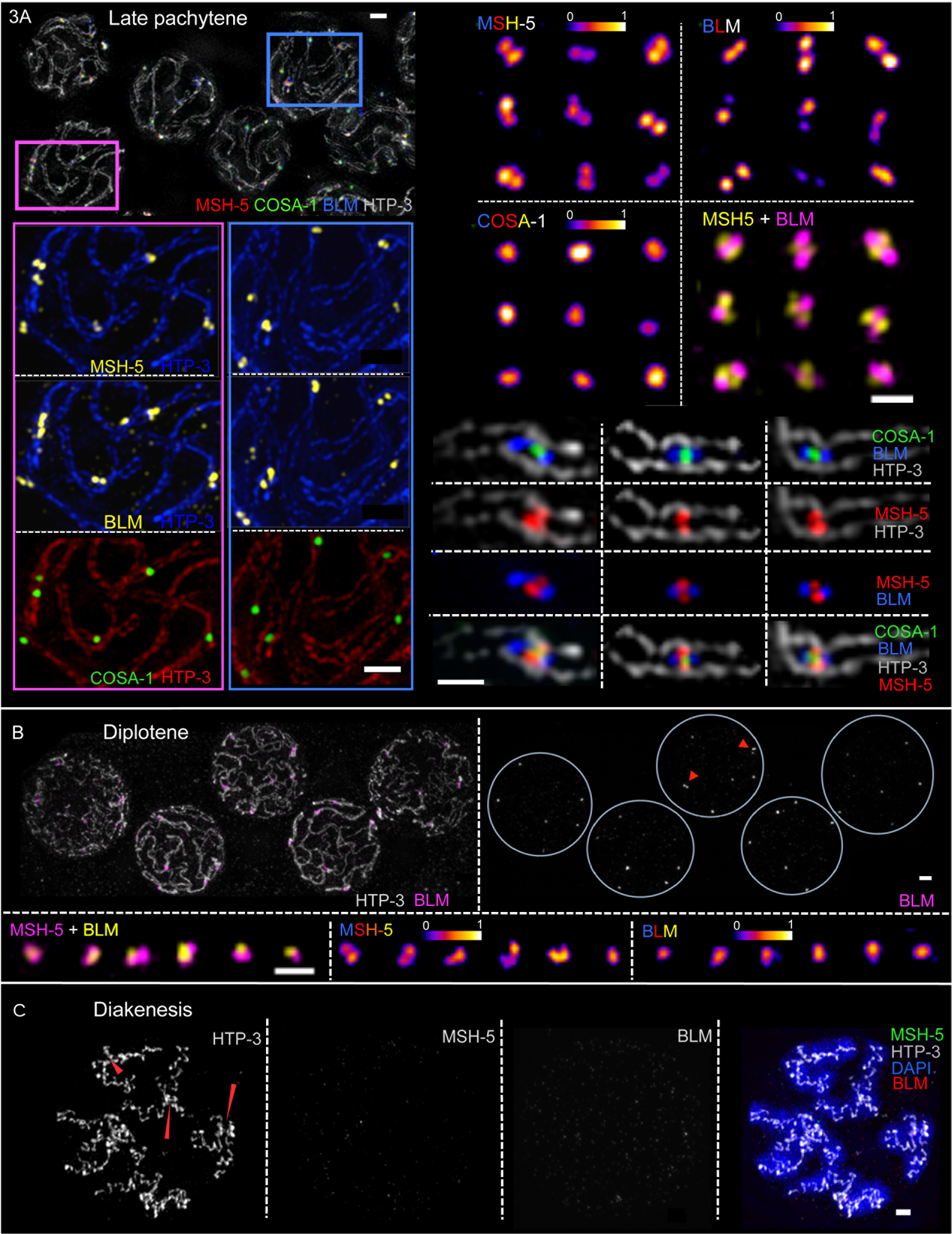
SIM images of CO-designated recombination sites during late prophase. **(A)** Topleft: Late pachytene nuclei from a spread gonad, probed for BLM::HA, MSH-5, GFP::COSA-1 and HTP-3. Bottom left: enlargements of portions of this field. Right: Individually cropped recombination sites from the field on the left. Examples displayed show CO-designated sites at chromosome segments imaged from a fully frontal aspect. Localization of two populations of BLM (oriented in parallel with the chromosome axes, marked by HTP-3), two populations of MSH-5 (oriented perpendicular to the axes) and a single population of COSA-1 at the center, together define five spatially resolvable domains within each CO-designated recombination site, located between the two homolog axes. The scale bars represent 1 μm (left panels) and 500 nm (right panels). **(B)** Top: Diplotene nuclei of a spread gonad, probed for BLM::HA and HTP-3. In contrast to late pachytene, most BLM signals are no longer reliably resolved as doublets at this stage; arrowheads highlight rare examples of BLM doublet foci at CO-designated sites in diplotene. Scale bar: 1 μm. Bottom: Individually cropped recombination sites from diplotene nuclei probed for BLM, MSH-5 and HTP-3. In contrast to BLM, MSH-5 signals retain a complex architecture at diplotene, similar to late pachytene. Scale bar: 500 nm. **(C)** Images of a diakinesis nucleus, indicating the presence of chiasmata, temporary connections between desynapsed homologs that reflect the formation of COs (red arrowheads); BLM and MSH-5 are no longer detected. Scale bar: 1 μm.

At these designated CO sites, the mean distance between the centroids of the paired BLM signals was 240 ± 34 nm (n = 20; Fig S3A) and the mean distance between paired MSH-5 signals was 168 ± 24 nm (n = 19; Fig S3A). Based on these measurements, we derive a lower-limit estimate of 290-300 nm, or about 800-900 bp, for the average length of the underlying DNA intermediates present at late pachytene CO-designated sites (see Methods and Discussion).

Further changes in CO site architecture accompany the transition to diplotene, when homologs start to desynapse and dissociate from each other. In contrast to pachytene, the majority of BLM signals can no longer be resolved as two foci, while two populations of MSH-5 are still detected at many CO designated sites during diplotene. The cohorts of proteins no longer exhibit any consistent arrangement relative to each other or to the chromosome axes (Fig 3B). Finally, by diakinesis, when the bivalents are completely de-synapsed, BLM and MSH-5 are no longer visible at the CO site, which is now cytological visible as a chiasma by using axis and DNA markers (Fig 3C).

### CO-designated recombination sites are encased by SC-CR proteins

Since meiotic recombination events progress and mature in the context of the SC, we used SIM imaging to investigate the spatial relationships between the SC-CR and the cohorts of DNA repair proteins present at recombination sites (Fig 4, S3A). When spreading is conducted using physiological or hypertonic conditions (“Hanks” in Fig S3A), SC destabilization is minimized (compared to hypotonic spreads, “H_2_O”) and a distance more similar to *in situ* spacing between the axes is maintained, reflecting better preservation of the SC-CR (Fig S3A-B). In early pachytene nuclei analyzed under such conditions, most signals for DNA repair proteins are found adjacent to, but not directly co-localizing with SC-CR proteins; *i.e.*, repair foci appear laterally associated with the SC-CR and only rarely appear embedded in it (Fig 4A and Fig S3B).

**Fig 4.**
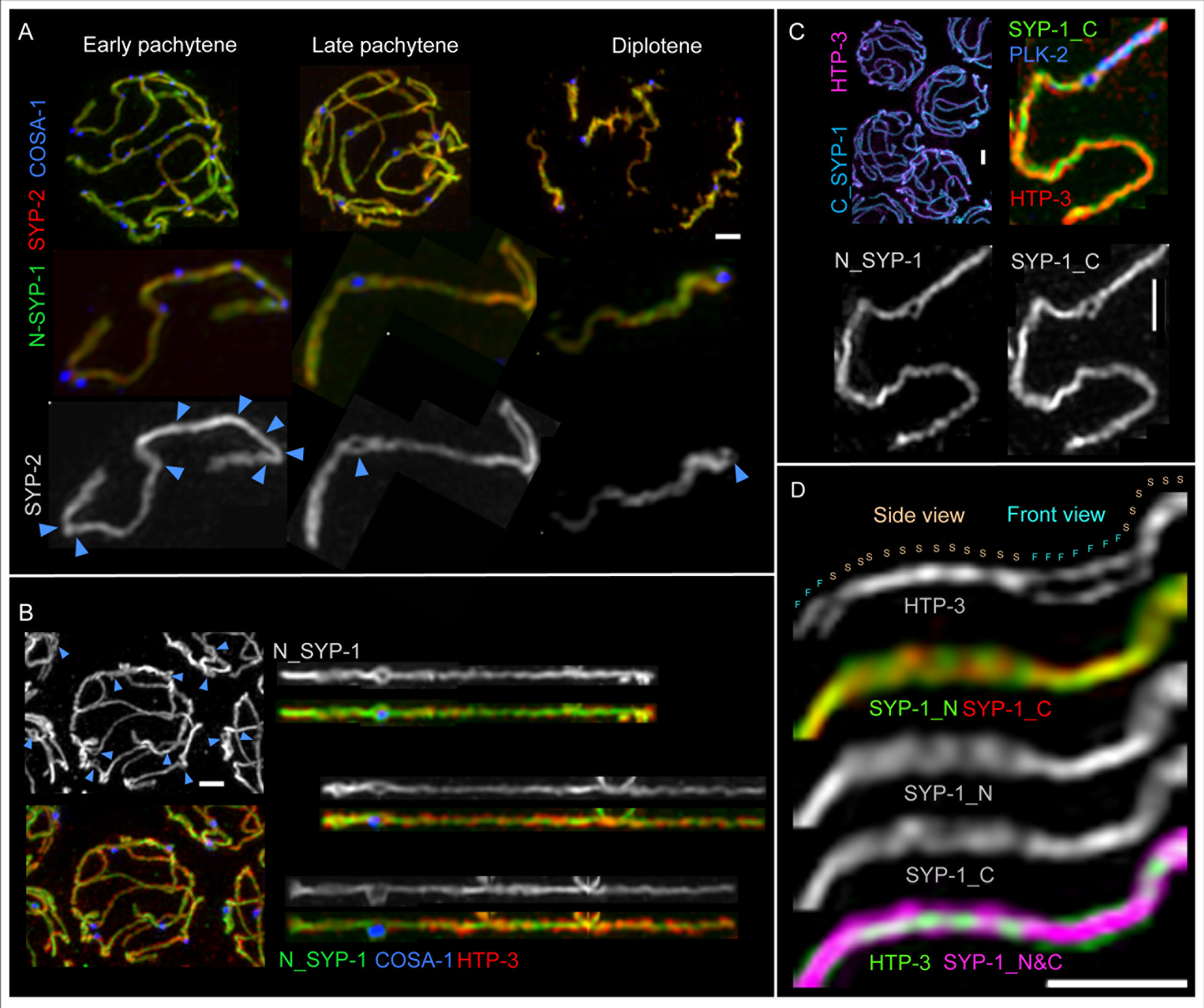
The SC-CR engulfs CO-designated sites as a bubble-like structure (see also Figure S3). **(A)** Top: Representative images of nuclei from early pachytene, late pachytene and diplotene stages, probed for GFP::COSA-1 (blue), SYP-1 (green) and SYP-2 (red). Bottom: one chromosome pair, individually cropped from each nucleus above. Blue arrowheads indicate the positions of multiple recombination sites in early pachytene and the single CO-designated recombination site at late prophase. **(B)** Representative late pachytene nucleus (left) and 3 individually straightened chromosome pairs (right) probed for GFP::COSA-1 (blue), SYP-1 (top: grey; bottom: green) and HTP-3 (red), illustrating that SC-CR bubbles are detected only at CO-designated sites (blue arrowheads). **(C)** Representative late pachytene nuclei (top left) and one individually-cropped SC (right) probed for PLK-2::HA, HTP-3, the N and the C-termini of SYP-1, illustrating that the SC-CR bubble is detected by antibodies against both the N- and C-termini of SYP-1. **(D)** Example of a segment of a late pachytene SC, illustrating both the frontal (F) and lateral (side; S) aspects of the SC. Scale bars represent 1 μm.

In contrast, during late pachytene, the SC-CR is consistently detected as a bubble-like structure at the CO-designated site, fully or partially surrounding the cohort of DNA repair proteins concentrated at the CO site (Fig 4A-C). A previous immuno-EM analysis (Schild-Prufert et al., 2011) showed that the SYP-1 N-terminus localizes at the center of the *C. elegans* SC, whereas the SYP-1 C-terminus localizes more laterally within the SC, closer to the chromosome axes, consistent with a head-to-head organization of SYP-1 within the SC-CR; the small SYP-2 protein localizes at the center of the SC. The SC-CR bubbles at CO-designated sites are detected with antibodies against both the N- and C-termini of SYP-1 and by antibodies against SYP-2. This bubble-like organization of the SC-CR around CO sites can be detected as early as differentiated CO sites first appear (*i.e.* in nuclei transitioning between early and late pachytene), and these CO-site SC bubbles are consistently detected at CO sites throughout the late pachytene stage (Fig 4A-C, Fig S3C). Encasement of recombination complexes by SC-CR proteins is even more pronounced in mutants where only a single chromosome pair in a nucleus harbors a recombination site, as this feature is exaggerated by the fact that SC-CR proteins become preferentially enriched on such chromosomes (Machovina et al., 2016; Nadarajan et al., 2017; Pattabiraman et al., 2017); Fig S3D).

Whereas previous work showed that polo-like-kinase PLK-2 is recruited to the SCs during pachytene (Harper et al., 2011; Labella et al., 2011; Pattabiraman et al., 2017), we note that PLK-2 is not part of the structure that encases the future CO site upon transition to late pachytene. Rather, PLK-2 is preferentially concentrated on the SC segment extending from the CO-designated site to the nearest chromosome end (referred to as the “short arm”), and also localizes within the SC-CR bubble together with the recombination proteins (Pattabiraman et al., 2017); Fig 4C, S3C). Further, while the CO-site SC-CR bubbles persist throughout late pachytene, they disappear concomitantly with disassembly of the SC. By mid-diplotene, remnants of the SC-CR bubble can be detected only on the side of the CO site adjacent to the short arm of the bivalent, where the SC-CR proteins remain specifically enriched at this stage (Nabeshima et al., 2005); Fig 4A).

Our SIM imaging of SC-CR proteins together with CO-site markers and chromosome axis marker HTP-3 (Goodyer et al., 2008; MacQueen et al., 2005) provides additional information regarding the structural organization of the *C. elegans* SC. As homologous chromosome pairs are twisted around each other during the pachytene stage, SC stretches can be visualized either in a frontal view, where the two chromosome axes (represented by HTP-3 immunostaining) are parallel to each other, or in a lateral view, in which HTP-3 appears as a single thin line (Fig 4D). When the SC is visualized in frontal views, SYP-2 and the N- and C-termini of SYP-1 all localize between the aligned chromosome axes, as expected. Interestingly, when the SC is visualized in lateral aspect, the SYP-1 N- and C-termini and SYP-2 are all detected on both sides of the single HTP-3 track, indicating the presence of SC-CR proteins both above and below the plane defined by the chromosome axes (Fig 4D).

Together, our data suggest that the *C. elegans* SC-CR is composed of (at least) two layers, similar to the 3D structures recently reported for the Drosophila and mouse SCs (Cahoon et al., 2017; Schucker et al., 2015), and the CO recombination intermediates become encased between these layers in a bubble-like structure, flanked on either side by the chromosome axis.

### SC-CR proteins play a role in promoting CO formation is distinct from the roles of other pro-CO factors

SC-CR proteins have been implicated in promoting CO formation in multiple organisms, and essentially all meiotic COs in *C. elegans* depend on the SC-CR proteins (MacQueen et al., 2002). However, the nature of the CO-promoting activit(ies) of the SC-CR is not well understood, and thus it has been unclear whether the SC-CR makes distinct contributions from other conserved pro-CO factors such as orthologs of MutSγ, ZHP-3 and COSA-1. Therefore, we analyzed and compared the dynamic localization of multiple DNA repair factors in two classes of mutants that fail to form COs: 1) null mutant that lack SC-CR proteins and thus fail to form SCs (*syp-1* and *syp-3*) (MacQueen et al., 2002; Smolikov et al., 2007); and 2) mutants that are proficient for SC assembly, DSB formation and DSB repair, but lack conserved pro-CO factors (*msh-4*, *cosa-1* and *zhp-3*, termed “recombination mutants” from here on; (Jantsch et al., 2004; Yokoo et al., 2012; Zalevsky et al., 1999).

In both SC-CR and recombination mutants, DSB repair sites marked by RAD-51 and/or RPA appear throughout early prophase and accumulate, as in WT. As early prophase is prolonged in these mutants (Rosu et al., 2013; Stamper et al., 2013; Woglar et al., 2013) the numbers of detectable DSB repair sites are higher and peak at a more proximal region of the gonad (Figure 5A). Similar to WT, BLM localizes to a subset of RPA sites during early prophase (Fig S4A). In both classes of mutants, MSH-5 foci can also be detected beginning in zygotene and accumulate until the transition to late pachytene (Figs 5B-C, S4B), albeit they appear fainter than in WT. Further, as in WT, these early MSH-5 foci appear to be largely dependent on the ability of chromosomes to engage in interactions with their homologs (Fig S5A).

However, in late prophase these two classes of mutants differ markedly from each other and from WT with respect to localization of pro-CO factors. In *cosa-1* and *zhp-3* mutants, MSH-5 foci are not stabilized, and they disappear concurrently with other DNA repair markers following transition to late pachytene (Fig 5B and Fig S4B). In contrast, in SC-CR mutants, MSH-5 foci can be detected following transition to a late pachytene-like state (Fig 5C and 5D). COSA-1 and ZHP-3 are also recruited to these late MSH-5-positive sites, and as in WT, RPA is lost from these late COSA-1/MSH-5/ZHP-3-marked sites (Fig 5D and Fig S5B). Further, as in WT, late prophase nuclei in SC-CR mutants consistently display no more than six foci (average of 4.2 foci per nucleus the *syp-3* mutant), and these foci are non-randomly distributed, such that two foci rarely occur on the same chromosome axis (p<0.0001). However, in contrast to WT, where BLM is maintained at CO-designated sites during late pachytene, BLM becomes undetectable at late COSA-1/MSH-5 marked sites soon after RPA is lost (Fig 5C).

**Fig 5.**
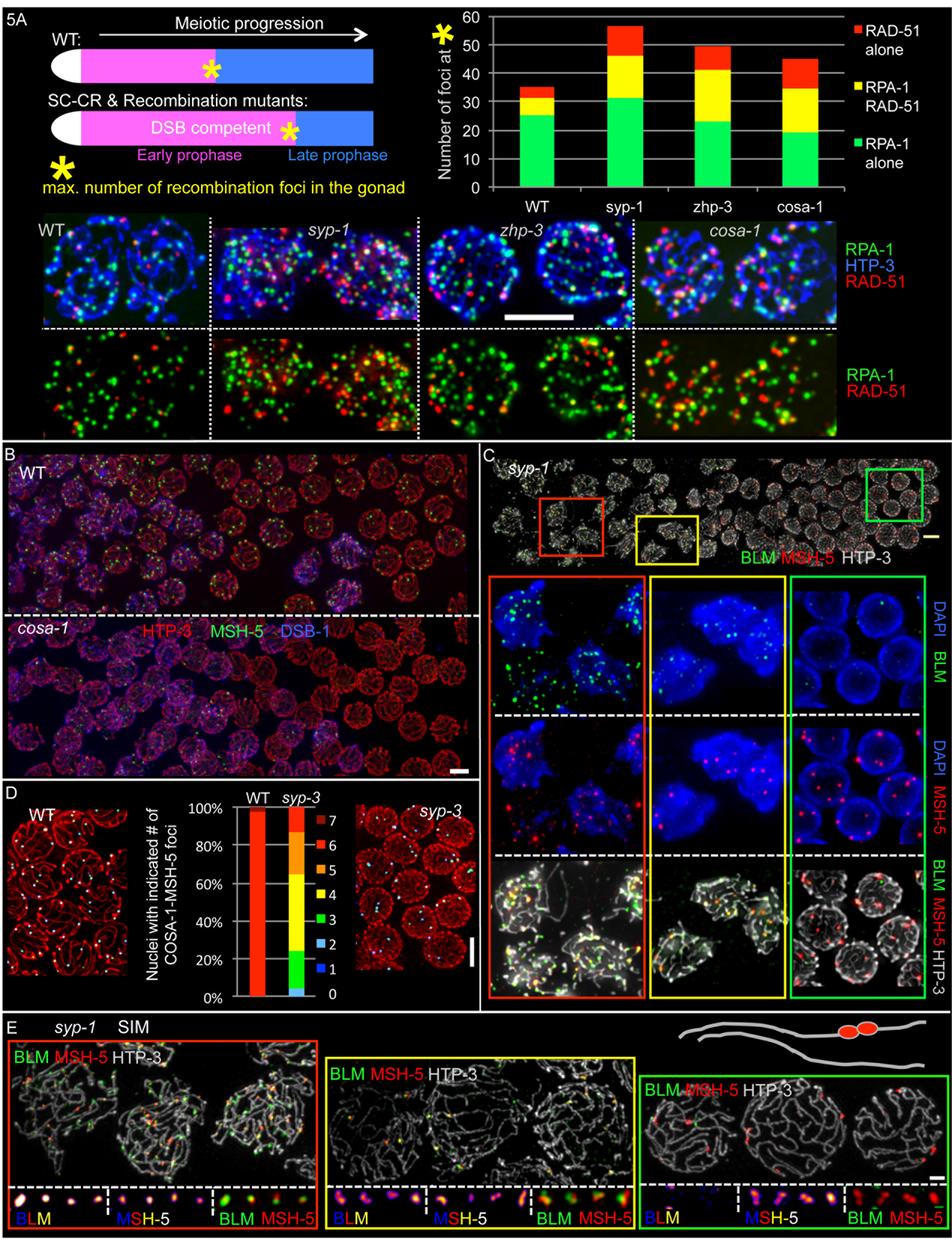
Dynamic localization of meiotic recombination factors during mutant meiosis (see also Figures S4 and S5) **(A)** Top left: Schematic depicting the positions of nuclei just prior to the early-to-late prophase transition (*), which have the highest numbers of recombination foci; as indicated, this transition is delayed in mutants deficient in crossing over. Top right: Quantification of RPA-1 and RAD-51 foci in spread nuclei from the positions indicated by the asterisk in the schematic; numbers of nuclei scored were: WT (n=34), *syp-1* (n=23), *cosa-1* (n=20) and *zhp-3* (n=20). For all three mutants, total numbers of recombination intermediates detected (RPA-1 and RAD-51 foci combined) were significantly higher in than in WT; P-values calculated using two-tailed heteroscedastic student's t-test were: *syp-1*, 2.5E-17; *cosa-1*, 1.7E-08; and *zhp-3:* 2.1E-05. Bottom: Representative examples of quantified nuclei from the early-to-late prophase transition, when numbers of recombination foci are highest. **(B)** Early-to-late prophase transition region of WT (top) and *cosa-1* mutant (bottom) gonad spreads, stained for DSB-1 (early prophase marker; blue), MSH-5 (green) and HTP-3 (red). MSH-5 foci are not retained after the transition to late prophase in the *cosa-1* mutant (see also Fig S4B). **(C)** Spread gonad of a *syp-1* mutant, which lacks the SC-CR, probed for BLM::HA (green), MSH-5 (red), HTP-3 (grey) and DAPI (blue). Individually cropped fields of nuclei from early prophase (red outline, left), early-to-late prophase transition (yellow outline, middle) and late prophase (green outline, right). Throughout the prolonged early prophase of *syp-1* mutants, MSH-5 colocalizes with BLM foci (left). At the transition to late prophase, MSH-5 foci reduce in number, and BLM is still present (middle). Upon transition to late prophase, BLM is lost from MSH-5-decorated sites (right). The scale bar represents 5 μm. **(D)** Images of late prophase nuclei from WT (*gfp::cosa-1*) and the *syp-3* mutant (*syp-3; gfp::cosa-1*) probed for HTP-3 (red) MSH-5 (blue) and GFP::COSA-1 (green), flanking a graph depicting quantification of MSH-5/COSA-1 foci in such nuclei (WT, n=35; *syp-3*, n=45). COSA-1 and MSH-5 do co-localize during late prophase in the *syp-3* mutant, but always to six or fewer sites per nucleus (average 4.2) **(E)** Top: representative SIM images of early prophase (red outline), early-to-late-prophase transition (yellow outline) and late prophase (green outline) nuclei from the *syp-1* mutant, probed for BLM::HA (green), MSH-5 (red), HTP-3 (grey). Scale bar represents 1 μm. In late prophase, 12 unpaired HTP-3 stretches can be observed. Bottom: enlarged individually cropped DNA repair sites from the same SIM fields; scale bar represents 500 nm. See text of Results for description of abnormal recombination site architecture in this mutant. Note that while MSH-5 is consistently detected as doublets in late prophase nuclei in the *syp-1* mutant, BLM is not associated with these sites, and the MSH-5 doublets are found in close association with and oriented along single unpaired axes, as indicated in schematic form above the late prophase images.

SIM imaging revealed that as in WT, MSH-5 signals in SC-CR mutants are detected as singlets in early prophase, and then can be resolved as doublets from the early-to-late prophase transition and throughout the remainder of the late pachytene like-stage. In contrast to WT, BLM is rarely detected as double foci in early prophase. At the early-to-late prophase transition, BLM is transiently detected as double foci associated with MSH-5-doublet marked sites; however, these structures do not consistently adopt the specific cruciform arrangement observed at CO-designated sites during WT meiosis, and they are not detected between parallel-aligned chromosome axes (Fig 5E). In late prophase when BLM can no longer be detected at MSH-5 marked sites, the paired MSH-5 foci in SC-CR mutants are both tightly associated with a single unpaired chromosome axis (Fig 5E). These structures may potentially reflect nonproductive remnants of earlier unstable IH repair intermediates, and/or they may represent the formation of CO-like recombination intermediates between sister chromatids rather than between homologs.

Thus, despite an inability to form IH COs, mutants lacking the SC-CR nevertheless are able to concentrate pro-CO factors at late prophase repair sites and generate recombination intermediates that share some features in common with the late pachytene CO-specific intermediates present during wild-type meiosis. However, these intermediates exhibit abnormal architecture and protein composition. Together the data indicate that SC-CR proteins play a role in promoting normal formation, architecture and stability of CO recombination complexes that is distinct from the roles of conserved CO-promoting factors such as COSA-1 and MutSγ that actually become concentrated at the CO sites.

## Discussion

### Simultaneous visualization of multiple recombination markers in spread germline nuclei illuminates the progression of meiotic recombination

Our work analyzing the localization of multiple recombination proteins together, in a context where signal-to-noise ratio has been greatly improved, has been highly informative in several ways:

First, our approach has allowed us to detect abundant RPA foci arising and accumulating during meiotic progression through the end of early pachytene. Moreover, based on timing, numbers, colocalization with BLM, and distinct localization relative to RAD-51, we can infer that the majority of these RPA foci represent markers of post-strand-exchange recombination intermediates. Second, analysis of MSH-5 localization in combination with RPA, RAD-51 and/or BLM, both in wild-type meiosis and in nuclei where specific chromosome segments cannot engage in homologous synapsis, indicates that MSH-5/RPA/BLM co-foci mark sites of IH recombinational repair. In further support of these inferences, chromosome segments that cannot engage in IH repair accumulate RAD-51 marked intermediates until the transition to late pachytene. Together, these observations and inferences strongly parallel the findings from cytological analyses of mouse meiosis in which localization of the same set of recombination complexes were analyzed in pairwise combinations (*e.g.* (Moens et al., 2002; Walpita et al., 1999). These striking parallels indicate that there is broad conservation of multiple early steps in the meiotic recombination program, despite differences regarding whether early steps in recombination are (as in mammals (Baudat et al., 2000; Edelmann et al., 1999; Romanienko and Camerini-Otero, 2000), or are not (as in *C. elegans*; (Dernburg et al., 1998) required to achieve synapsis between the homologous chromosomes. Thus, insights gained from simultaneous analyses of multiple recombination factors in the *C. elegans* system will be relevant for extending our understanding of mammalian reproduction.

Third, simultaneous immunolocalization of specific combinations of three recombination proteins (*i.e.* RAD-51 and RPA together with either BLM, MSH-5 or COSA-1) enables improved accounting of the total number of recombination events present during wild-type meiosis. We find that by the end of the early pachytene stage, after DSBs have been introduced and converted into repair intermediates over many hours, the total number of active repair sites detected using such three-marker combinations peaks at 25-40 per nucleus before declining upon transition to late pachytene. Accordingly, we conclude that each of the six chromosome pairs in a nucleus receives an average of at least 4-7 breaks over the course of meiotic prophase. These numbers may closely approximate the actual numbers of DSBs introduced, since the peak number of total repair foci detected using this three-marker approach correlates with the duration of early prophase (*i.e.* when comparing WT and meiotic mutants where early prophase is prolonged). Further, this estimate of DSB number is compatible with the average number of γ-ray generated DSBs (3.9 per chromosome pair) required to reliably restore COSA-1 foci in the absence of endogenous meiotic DSBs (Yokoo et al., 2012).

Based on our observations, we propose that the aforementioned three-marker combinations enable visualization of virtually all ongoing meiotic recombination events present in a given nucleus at a given moment in time. Accordingly, we hypothesize that DSB-dependent recombination intermediates generated during early prophase accumulate until the transition to the late pachytene stage, but repair is not completed until after this transition. Further, as recombination markers disappear from NCO sites following transition to late pachytene while being retained at CO sites, we infer that completion of NCO repair likely occurs earlier than completion of CO repair. Differential timing for completion of NCO and CO repair is reminiscent of the differential timing of appearance of NCO and CO products demonstrated to occur in budding yeast meiosis (Allers and Lichten, 2001). However, in contrast to the situation in budding yeast, where NCOs (but not COs) can be generated prior to a programmed transition to late pachytene, in *C. elegans*, completion of both NCO and CO events appears to be dependent on the analogous development transition.

### Inferring the architecture of underlying recombination intermediates

SIM images of recombination proteins at DSB repair sites enable us to make inferences and hypotheses regarding the types of recombination intermediates that may be present, their relative timing of appearance and key transitions in the process.

First, our finding that RAD-51 foci usually occur as doublets indicates that following DSB formation, both DNA ends undergo resection and loading of DNA strand-exchange proteins. Coupled with the analogous finding that strand-exchange proteins Dmc1p and Rad51p co-occupy both ends of a DSB during *S. cerevisiae* meiosis (Brown et al., 2015), we infer that dual end resection is a conserved feature of the meiotic program and that both DSB ends may have the potential to participate in homology search.

Second, we infer that resected DSBs rapidly give way to post-strand-exchange recombination intermediates. Such intermediates accumulate throughout early prophase and are typically decorated by two populations of BLM and two populations of RPA, frequently flanking a single population of MSH-5; notably, however RAD-51 is not detected at most of these sites. Given the initial presence of RAD-51 on both DSB ends, its absence from the majority these inferred post-strand-exchange intermediates implies that the behavior of the two DSB ends must be coupled. This may involve temporally coupled invasion of the same DNA duplex by both ends and/or rapid second-end capture following initial strand invasion by one end. Alternatively, the two DSB ends might be coordinated in a more complex, regulatory manner; *e.g.* strand invasion by the first end could be coupled to RAD-51 filament removal and replacement by RPA on the ssDNA tail of the second end, as has been proposed to occur during synthesis-dependent strand annealing (SSDA) mediated DSB repair in *S. cerevisiae* (Liu et al., 2017; Mitchel et al., 2013). Such a mechanism could prevent the second end from engaging a different chromatid while also facilitating eventual second end capture.

Changes in organization of recombination sites that occur following the transition to the late pachytene stage presumably reflect changes in the underlying structure of meiotic recombination intermediates. We propose that the transition from one to two populations of MSH-5 and the loss of RPA at CO-designated recombination sites reflects conversion of earlier intermediates to CO-specific intermediates that lack ssDNA stretches, reflecting completion of repair DNA synthesis at those sites. We note that the estimated lengths of the underlying DNA intermediates present at these sites (800-900 bp; see Methods) is compatible with the size scale of CO recombination intermediates deduced from average lengths of CO-associated gene conversion tracts in budding yeast (1.6 – 2 kb) and mammals (500-600 bp) (Cole et al., 2014; Mancera et al., 2008; Martini et al., 2011). Moreover, the organization of the recombination factors present at these sites is consistent with the interpretation that these likely represent double Holliday junction (dHJ) recombination intermediates, which have been demonstrated to be CO-specific recombination intermediates during yeast meiosis (Schwacha and Kleckner, 1995). Accordingly, we envision that the two population of MSH-5 represent two cohorts of MutSγ sliding clamp complexes (Snowden et al., 2004) that embrace and accumulate on each of the two parallel DNA duplexes that span the distance between the two junctions, and that the two populations of BLM helicase complexes localize near or at the junctions themselves (Jagut et al., 2016). In this proposed organization, the MutSγ complexes could serve as a roadblock that would serve as an impediment to branch migration, thereby inhibiting conversion of such intermediates into NCO products via dHJ dissolution.

A remarkable feature of the meiotic program is that eventual resolution of CO-specific recombination intermediates must occur in a highly biased manner so that each CO-designated intermediate matures in a manner that reliably yields CO (rather than NCO) products. Models in which dHJs are the relevant CO intermediates require that the two junctions consistently be resolved in “opposite sense” to generate a CO (Szostak et al., 1983), but how such biased resolution might be accomplished is completely mysterious. One possibility is that pre-existing asymmetry in the DNA of the intermediate, *e.g.* newly synthesized DNA, or nicks (if the HJs are not ligated), could be used as a signal to direct cutting by resolvases. Alternatively, architecture of the recombination intermediate in the context of meiotic chromosome structure could potentially provide the information needed to achieve biased resolution. We observed a change in the organization of CO recombination sites during meiotic progression that may provide insight this issue: in contrast to pachytene, BLM foci are no longer detected as doublets at diplotene CO sites. Based on similarity to the timing of appearance of CO products during budding yeast meiosis (Allers and Lichten, 2001), this change in CO-site organization may be coupled to resolution of CO intermediates. One possible explanation for this change is that dHJ resolution may occur in two temporally distinct steps, with CO biased resolution resulting from chromosomal features that become highly asymmetric at this stage (see below). Another possible scenario is that large-scale reorganization of chromosomes at the diplotene stage enables a 3D reconfiguring of dHJ intermediates that results in the two HJs being brought into close spatial proximity, thereby enabling coupled resolution.

### Structure of the SC-CR and roles in promoting the formation of meiotic COs

Our imaging of the SC-CR in relation to recombination sites, in combination with our analysis of recombination sites in *syp* mutants, provides new insight into the potential functions of the SC-CR in promoting the formation of meiotic COs.

First, our finding that late prophase MSH-5/COSA-1 marked sites appear non-randomly distributed on the 12 now unpaired chromosome axes in *syp* mutants suggests that *C. elegans* chromosomes retain a capacity to communicate along, between and/or among each other to limit the number of “CO-like” sites among a surplus of precursors even when the SC-CR is absent, similar to findings previously reported for CO site markers during yeast meiosis (Fung et al., 2004). However, *syp* mutants display significantly fewer “CO-like” sites in late prophase than the six designated CO sites consistently observed during wild-type meiosis. Thus, we envision that the previously-described CO-limiting function of the SYP proteins (Hayashi et al., 2010; Libuda et al., 2013) is needed to counteract a strong impetus promoting CO designation and maturation that operates when the SYP proteins are present.

Second, our data argue for a role for the SC-CR in creating spatially-segregated compartments that could enable distinct biochemistry to yield different outcomes of repair at CO and NCO sites within the same nucleus. SIM imaging revealed that upon transition to late pachytene, the cohorts of recombination proteins that accumulate at CO-designated sites, and the underlying recombination intermediates, become encased by SC “bubbles”. At the sites of these bubbles, CO-designated recombination events appear sandwiched between two layers of SC-CR proteins and embraced laterally by axial structures that are assembled along the length of each homolog pair. This organization effectively creates two distinct compartments, *i.e.*, inside and outside the bubble. We propose that the “inside” environment favors local recruitment and retention of pro-CO factors that promote maturation of CO intermediates. Further, the inside environment would simultaneously protect intermediates from activities that dismantle recombination intermediates. Conversely, recombination intermediates located in the outside environment would be susceptible to factors that dissociate recombination intermediates and complete repair *via* mechanisms that generate NCO products, *e.g.* SDSA, dHJ dissolution and/or single HJ cleavage followed by branch migration. Basically, we propose that by segregating CO and NCO intermediates into spatially distinct compartments, the SC-CR can protect the CO sites from NCO-promoting activities exerted at the same time at all other repair sites within the same cell.

This idea that the SC-CR functions in preventing CO-designated intermediates from becoming dismantled prematurely is reinforced by our analysis of recombination sites structure in mutants lacking SC-CR components. Some structures that bear resemblance to CO-designated sites in the WT do form in these mutants, but these sites lose BLM prematurely, concurrently with the disappearance of DNA repair markers from all other repair sites.

SIM imaging of the organization of CO recombination sites relative to SC-CR components at the diplotene stage leads us to speculate that the SC-CR might also play an additional, late, pro-CO role, *i.e.* in promoting CO-biased resolution of dHJs. We noted above that CO sites undergo a change in organization during diplotene that may correspond to the timing of resolution. Remnants of the SC-CR bubble are specifically retained at only one side of the CO site at this stage, raising the possibility that presence of SC-CR proteins in close proximity to one junction and absence from the other junction could play a role in determining distinct resolution outcomes.

## Author contributions

A.W. designed and conducted the experiments, A.W and AM.V. analyzed the data and wrote the manuscript.

## Acknowledgments

We thank the Caenorhabditis Genetics Center (funded by NIH Office of Research Infrastructure Programs P40 OD010440), M. Colaiacovo, A. Dernburg, A. Garnter, Y. Kim, D. Libuda, and R. Lin for strains and antibodies. We are grateful to V. Jantsch for allowing usage of the *him-6::HA* transgene prior to publication, to J. Loidl for sharing Lipsol, to F. Klein and J. Loidl for sharing experience in cytological preparations, to J. Mulholland for exterminating all the bugs in the microscope and to A. MacQueen for inspiration. We thank C. Akerib, C. Girard, V. Jantsch, and B. Roelens for critical reading of the manuscript. This work was supported by an FWF Erwin Schrödinger Fellowship (J-3676) to AW, by NIH grants R01GM53804 and R01GM67268 to AMV, by a pilot project under NIH/NICHD center grant U54 HD068158, and by American Cancer Society Research Professor Award RP-15-209-01-DDC to AMV. The project was also supported in part by Award Number 1S10OD01227601 from the National Center for Research Resources (NCRR). Its contents are solely the responsibility of the authors and do not necessarily represent the official views of the NCRR or the National Institutes of Health. The funders had no role in study design, data collection and analysis, decision to publish, or preparation of the manuscript.

## Methods

### Strains used in this study

- N2
- AV776: *spo-11(me44)* IV / *nT1[qIs51 let-?]* (IV;V)
- AV596: *cosa-1(tm3298)* / *qC1[qIs26]* III
- AV115: *msh-5(me23)* IV / *nT1[unc-?(n754) let-?]* (IV;V)
- AV818: *meIs8[unc-119(+); Ppie-1::gfp::cosa-1]* II; *cosa-1(tm3298)* III
- AV954: *him-6(jf93[him-6::HA]* IV
- AV831: *meIs8[unc-119(+); Ppie-1::gfp::cosa-1]* II; *cosa-1(tm3298)* III; *spo-11(me44) / nT1[qIs51 let-?]* IV;V
- CA258: *zim-2(tm574)* IV
- CA151: *him-8(me4)* IV
- AV307: *syp-1(me17)* V / *nT1[unc-?(n754) let-?]* (IV;V)
- AV842: *meIs8[unc-119(+); Ppie-1::gfp::cosa-1]* II; *cosa-1(tm3298)* III; *him-6(jf93[him-6::HA]* IV
- WS458: *unc-119(ed3)* III; *opls 263[rpa-1p::rpa-1::YFP + unc-119]*
- AV687: *syp-3(ok758)* I / *hT2[bli-4(e937) let-?(q782 qls48]* (I;III); *meIs8[unc-119(+); Ppie-1::gfp::cosa-1]* II
- CV2: *syp-3(ok758) I* / *hT2[bli-4(e937) let-?(q782 qls48]* (I;III)
- VC1474: *top-2(ok1930) /mIn1[mls14 dpy-10(e128)]* II (for experiments described VC1476 was mated with N2 males and F1 worms heterozygous for *mIn1* were analyzed)
- UV116: *him-6(jf93[him-6::HA]* IV
- AV959: *him-6(jf93[him-6::HA]* IV; *syp-1(me17)* V / *nT1[unc-?(n754) let-?]* (IV;V)
- AV960: *cosa-1(tm3298)* / *qC1[qIs26]* III; *him-6(jf93[him-6::HA]* IV
- AV961: *zhp-3(me95)* I / *hT2[bli-4(e937) let-?(q782 qls48]* (I;III); *him-6(jf93[him-6::HA]* IV
- AV962: *him-14(it44); him-6(jf93[him-6::HA]* IV
- AV963: *him-8(me4)* IV; *syp-1(me17)* V / *nT1[unc-?(n754) let-?]*

### Cytological preparations

Spreading of *C. elegans* gonads was performed as in (Pattabiraman et al., 2017). The gonads of 20–100 adult worms were dissected in 5μl dissection solution (see below) on an EtOH-washed 22×40mm coverslip. 50μl of spreading solution (see below) were added and gonads were immediately distributed over the whole coverslip using a pipette tip. Coverslips were left to dry at room temperature (approximately 1 hour) and post-dried for two more hours at 37°C, washed for 20 minutes in methanol at −20°C and rehydrated by washing 3 times for 5 minutes in PBS-T. A 20-minute blocking in 1% w/v BSA in PBS-T at room temperature was followed by overnight incubation with primary antibodies at 4°C (antibodies diluted in: 1% w/v BSA in PBS-T supplied with 0.05% w/v NaN_3_). Coverslips were washed 3 times for 5 minutes in PBS-T before secondary antibody incubation for 2 hours at room temperature. After PBS-T washes, the nuclei were immersed in Vectashield (Vector) and the coverslip was mounted on a slide and sealed with nail polish. Dissection solution: For most experiments, gonads were dissected in 0.1% v/v Tween-20 in H_2_0. For nearly all experiments in which the SC-CR was visualized, gonads were dissected in 85% v/v Hank’s Balanced Salt Solution (HBSS, Life Technology, 24020-117) with 0.1% v/v Tween-20; the exceptions were for images shown in Figures S1C and S3D, where gonads were dissected in H_2_O as above. Results similar to dissection in HBSS were obtained by dissecting gonads in increasing concentrations of PBS, Dulbecco’s Modified Eagle’s Medium (DMEM, Sigma-Aldrich, D6546), or similar (data not shown). Spreading solution: (for one coverslip, 50μl): 32μl of Fixative (4% w/v Paraformaldehyde and 3.2–3.6% w/v Sucrose in water), 16μl of Lipsol solution (1% v/v/ Lipsol in water), 2μl of Sarcosyl solution (1% w/v of Sarcosyl in water).

### Antibodies used in this study

The following primary antibodies used: Chicken anti-HTP-3 (1:500) (MacQueen et al., 2005), guinea pig anti-HTP-3 (1:500) (MacQueen et al., 2005), rabbit anti-MSH-5 (1:10000) (SDI/Novus), chicken anti-GFP (1:2000) (Abcam), mouse anti-GFP (1:500) (Roche), rabbit anti-GFP (1:500) (Yokoo et al., 2012), mouse anti-HA (1:1000) (Covance), guinea pig anti-ZHP-3 (1:500) (Bhalla et al., 2008), rabbit anti-PLK-2 (Nishi et al., 2008), guinea pig anti-SYP-1 (1:200) (MacQueen et al., 2002), rat anti-RAD-51 (1:200) (Schvarzstein et al., 2014), rabbit anti-RPA-1 (1:500) (Lee et al., 2010), mouse anti-H3K4me2 (1:2000) (Cell Signaling Technology), rabbit anti-SYP-1 (1:2000) (MacQueen et al., 2002) and rabbit anti-SYP-2 (1:200) (Colaiacovo et al., 2003). Secondary antibodies conjugated to Alexa dyes 405, 488, 555 or 647, obtained from Molecular Probes, were used at 1:500 dilution. In cases where antibodies raised in mouse and guinea pig were used on the same sample, we used highly cross-absorbed goat anti-mouse secondary antibodies, obtained from Biotum (conjugated to CF488, or CF555 respectively) for secondary detection of the mouse primary antibody in order to avoid cross-reaction against antibodies raised in guinea pig.

### Imaging

Imaging, Deconvolution and 3D-SIM reconstruction were performed as in (Pattabiraman et al., 2017). Spreading results in squashing of *C. elegans* germline nuclei from 5 to 1-2 μm in thickness. Wide field (WF) images were obtained as 200 nm spaced Z-stacks, using a 100x NA 1.40 objective on a DeltaVison OMX Blaze microscopy system, deconvolved and corrected for registration using SoftWoRx. Subsequently, gonads were assembled using the “Grid/Collection” plugin (Preibisch et al., 2009) in ImageJ. For display, assembled gonads were projected using maximum intensity projection in ImageJ. 3D-Structured Illumination microscopy images were obtained as 125 nm spaced Z-stacks, using a 100x NA 1.40 objective on a DeltaVison OMX Blaze microscopy system, 3D-reconstructed and corrected for registration using SoftWoRx. For display, images were projected using maximum intensity projection in ImageJ or SoftWoRx. For display, contrast and brightness were adjusted in individual color channels using ImageJ. For Figure 4B and Figure S3C, SCs were computationally straightened using ImageJ 2D-straightening tool on well-spread chromosomes (250-1250 nm maximum Z-stack thickness).

### Quantification of recombination foci

To quantitatively display the appearance, disappearance and co-localization of recombination foci in a single spread gonad (as *e.g.* in Fig 1), all nuclei within the gonad that did not overlap with other nuclei were considered (usually 6-12 nuclei per cell row). Meiotic entry was defined by the appearance of HTP-3 as continuous tracks along chromosomes in the majority of nuclei in the cell row. Recombination foci were counted manually on deconvolved and stitched images. We considered a signal to be a recombination focus when found within a nucleus, colocalizing with HTP-3 and 5 times brighter than background.

### Measurement/estimation of distances between meiotic chromosome axes and paired recombination foci

The centroids of a recombination focus, or axial signals, respectively were determined as the brightest pixel(s) within the signal (in raw data, one pixel in XY represents 80.1 × 80.1 nm). As chromosome pairs twist around their longitudinal axes, the fullest extent of separation between individual HTP-3 signals (in the XY plane) is observed when the SC is visualized in a fully frontal aspect. Therefore, to determine the separation between HTP-3-marked chromosome axes, we measured the distances at 7-9 different sites along a pair of aligned homologous chromosomes in segments that displayed separation of HTP-3 axis signals, with the three highest values being recorded. The distances between the centroids of paired recombination foci (MSH-5-MSH-5 or BLM-BLM) were measured (in nuclei spread using hypotonic conditions) at sites where parallel axes were visualized in single XY plane in a fully frontal view, as indicated by at least 190 nm of axial separation.

### Estimation of DNA length underlying late recombination structures

We used the Pythagorean theorem to estimate the length of the DNA underlying late pachytene cohorts of recombination factors based on the mean measured MSH-5-MSH-5 (m = 168) and BLM-BLM (b =240) distances, using the formula: d = 2[ (m/2)^2^ + (b/2)^2^]^1/2^, yielding a value of 293 nm. As a dHJ or similar recombination intermediate of this length is expected to be composed primarily of B-form DNA, and as 1 kb of B-form DNA has a length of 340 nm, we estimate that the average length of DNA underlying late pachytene CO-site structures is on the order of 800-900 bp.

## Supplemental Information

### Supplemental Figure Legends

**Fig S1(related to Figure 1).**
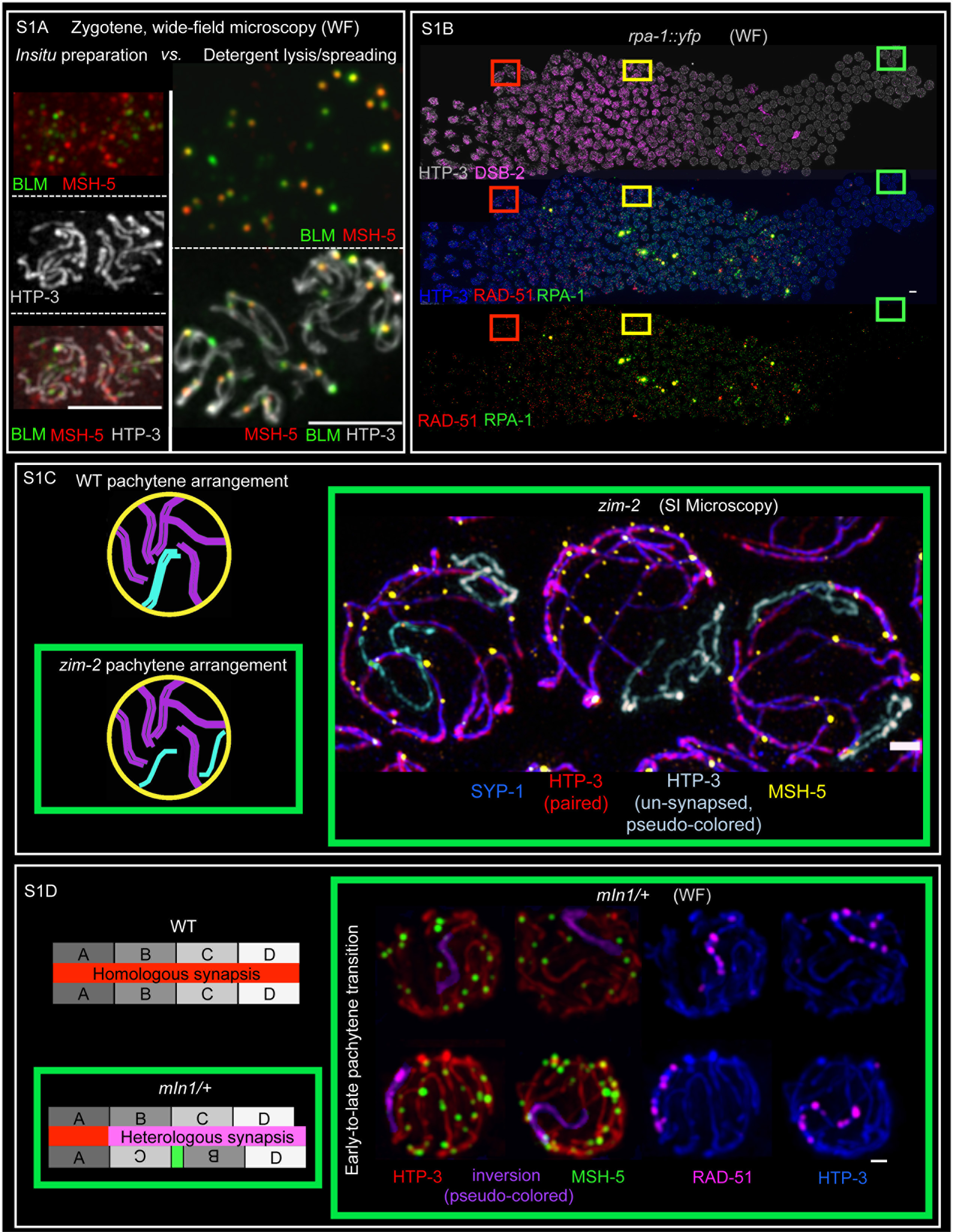
(A) Nuclear lysis and chromosome spreading improve visualization of DNA repair complexes at sites of ongoing meiotic recombination. Example images of zygotene/early pachytene nuclei prepared using either a whole mount *in situ* method (according to: (Jagut et al., 2016); left panel) or detergent-based nuclear lysis and chromosome spreading procedure (according to: (Pattabiraman et al., 2017); right panel), probed for: BLM, a DNA helicase [HIM-6 tagged by HA (*him-6::HA*), expressed from the endogenous locus; green], MSH-5, a pro-crossover factor (red) and HTP-3, a core component of the meiotic chromosome axis (grey). Wide-field microscopy with subsequent 3D-deconvolution and Z-stack projection is indicated as “WF”; scale bars represent 5 μm. **(B) Appearance, disappearance and localization of RAD-51 and RPA-1 and over the course of meiotic prophase.** Spread WT gonad probed for RAD-51 (red), RPA-1 (green), DSB-2 (marker of early prophase; purple) and HTP-3 (grey in the top panel and blue in the middle panel). Meiotic prophase progresses from left (leptotene) to right (diplotene). Boxes represent the enlarged fields displayed in Fig 1A. Scale bar represents 5 μm. **(C-D)) MSH-5 marks inter-homolog DNA repair events. (C)** Left: Schematics of pachytene chromosome arrangement in WT and in *zim-2* mutants, which are specifically defective for pairing and synapsis of chromosome V (cyan). Right: SIM images of spread nuclei from *zim-2* gonads at the early-to-late prophase transition, probed for: MSH-5 (yellow), SYP-1 (blue) and HTP-3 (red, if localizing with SYP-1, and pseudo-colored cyan when unsynapsed, indicating the unpaired Chr. V). MSH-5 foci are found on all synapsed chromosome pairs but not on the two unsynapsed chromosomes. Scale bar represents 1 μm. **(D)** Left: Schematics comparing WT chromosome II synapsis configuration with the chromosome II synapsis configuration in worms heterozygous for *mIn1*, which carries a large internal inversion and an integrated high-copy transgene array (indicated in green). Right: wide-field images of spread nuclei from gonads of *mIn1*/+ animals, probed for RAD-51 (magenta), MSH-5 (green), HTP-3 (red) and SYP-1 (blue). MSH-5 foci are absent from the chromosomes segments that are enriched for RAD-51-marked intermediates, which are presumed to correspond to heterosynapsed chromosome segments where interhomolog recombination intermediates cannot form; these segments are pseudo-colored in purple in the left panels. Scale bars represent 5 μm.

**Fig S2 (related to Figure 1).**
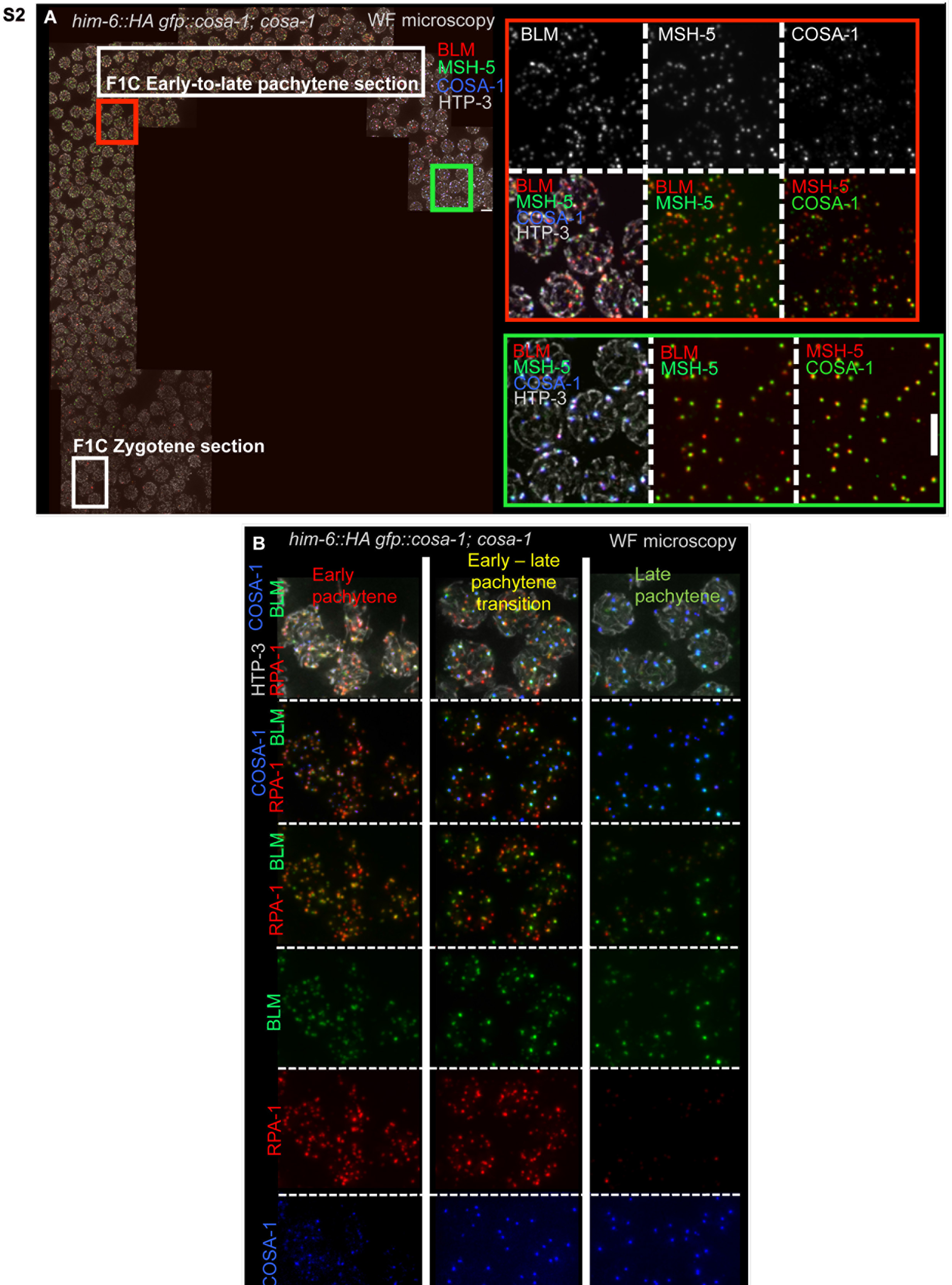
Dynamic localization of BLM, MSH-5, COSA-1 and RPA-1 during meiotic prophase. **(A)** Spread gonad, probed for BLM (HIM-6::HA, red), MSH-5 (green), GFP::COSA-1 (blue), and HTP-3 (grey). COSA-1 is tagged with GFP (*gfp::cosa-1*) and expressed from an randomly integrated overexpression construct in the background of the *cosa-1(tm3298)* deletion. Representative fields of early pachytene, early-to-late pachytene transition and late pachytene demonstrate that all COSA-1 foci localize with MSH-5 foci, and MSH-5 foci nearly always colocalize with BLM foci. BLM, in contrast, can be detected without MSH-5/COSA-1, especially during the early-to-late prophase transition. Please note changes of colors for different markers in the enlargements to visualize co-localization. Scale bars represent 5 μm. **(B)** Representative fields of early pachytene, early-to-late pachytene transition and late pachytene nuclei, probed for GFP::COSA-1 (blue), RPA-1 (red), BLM (HIM-6::HA, green) and HTP-3 (grey). During early pachytene, BLM foci strictly colocalize with RPA-1 foci. At the early-to-late pachytene transition, BLM and RPA are frequently found at different sites. In late pachytene and diplotene, BLM foci at designated CO sites, which are co-decorated by a bright COSA-1 focus, are largely free of RPA-1. Scale bars represent 5 μm.

**Fig S3 (related to Figures 3 and 4).**
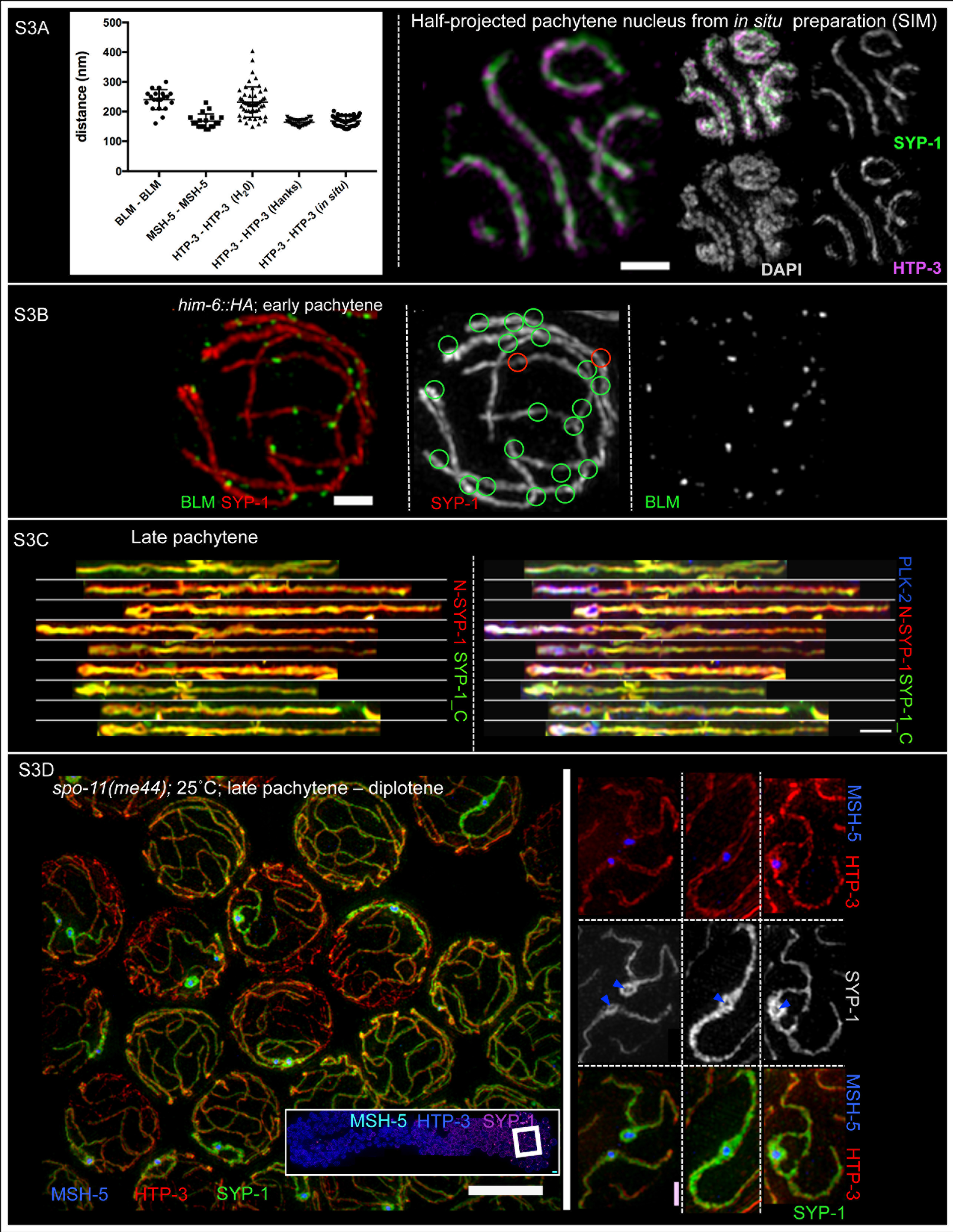
**(A)** Left, graph indicating distances measured between paired BLM foci and paired MSH-5 foci at late pachytene CO-designated sites (as depicted in Figure 3) and distances separating paired chromosome axes (as indicated by HTP-3 immunostaining) measured in different types of cytological preparations. The SC-CR is partially destabilized under the hypotonic spreading conditions (“H_2_O”) used here for visualization of recombination site architecture in the context of chromosome axes (Pattabiraman et al 2017), resulting in a significant increase (up to two-fold) in the separation between homologous chromosome axes in late pachytene nuclei relative to the roughly 120-170 nm distances measured *in situ* (Kohler et al., 2017; Schild-Prufert et al., 2011); this work)). Under the more physiological/hypertonic salt conditions used for visualizing recombination sites in relation to the SC-CR (“Hanks”; see Methods), the SC-CR is more stable, and inter-axis distance measurements are indistinguishable from measurements made using SIM images of SCs in nuclei from whole mount *in situ* preparations. Horizontal lines indicate the mean, error bars indicate standard deviation. Numbers of measurements represented in the graph (n) were as follows: BLM-BLM, n=20; MSH-5-MSH-5, n=19; HTP-3-HTP-3 (H_2_O), n=52; HTP-3-HTP-3 (Hanks), n=30; HTP-3-HTP-3 (*in situ),* n=51. Right, sample SIM images of SCs from a whole-mount *in situ* preparation; scale bar: 1 μm. **(B)** Example SIM image of an early pachytene nucleus, probed for the SC-CR protein SYP-1 (red) and BLM (green). In most cases, BLM appears laterally associated with continuous stretches of SC-CR proteins (indicated by green circles). At only few sites, BLM directly co-localizes with the SC-CR-protein (red circles). The scale bar represents 1 μm. **(C)** Examples of computationally straightened late pachytene SCs probed for the N-terminus (red) and C-terminus (green) of SYP-1. SYP-1 forms a continuous stretch along the bivalent, and at only one site per bivalent, a clear SYP-1 bubble is visible. Straightened chromosomes were aligned at the bubble. Right: PLK-2::HA (blue), a Polo-like kinase, localizes from late pachytene onward to the short arm of the bivalent, *i.e.* from the CO site to the nearest chromosome end; PLK-2::HA also forms a focus inside the SC-CR bubble. The scale bar represents 1 μm. **(D)** Field of late pachytene/diplotene nuclei from a *spo-11* gonad, which lacks the enzyme responsible for forming canonical meiotic DSBs, probed for HTP-3 (red), SYP-1 (green) and MSH-5 (blue). Although *spo-11* mutants are unable to generate canonical meiotic DSBs, a subset of chromosomes (5% at 25°C) nevertheless acquires a single recombination focus during late prophase, presumably reflecting the occurrence of spontaneous DNA lesions that recruit meiotic DNA repair proteins. The SC-CR proteins become preferentially concentrated specifically on the subset of chromosomes that received a recombination focus (Machovina et al., 2016; Nadarajan et al., 2017; Pattabiraman et al., 2017), and on such chromosomes, the SC-CR becomes prominently enriched around and engulfs these recombination foci. These foci most likely do not mark *bona fide* recombination events, as: 1) These sites do not reliably adopt the complex architecture observed at meiotic recombination sites in the WT; 2) These sites appear in many cases to occur between sister chromatids, accompanied by local alterations in synaptic configuration never seen in the WT (*e.g.* example on the right); crossover recombination is absent in *spo-11* when measured genetically (Dernburg et al., 1998). Scale bars represent 5 μm (large field) and 1 μm (inset).

**Fig S4 (related to Figure 5).**
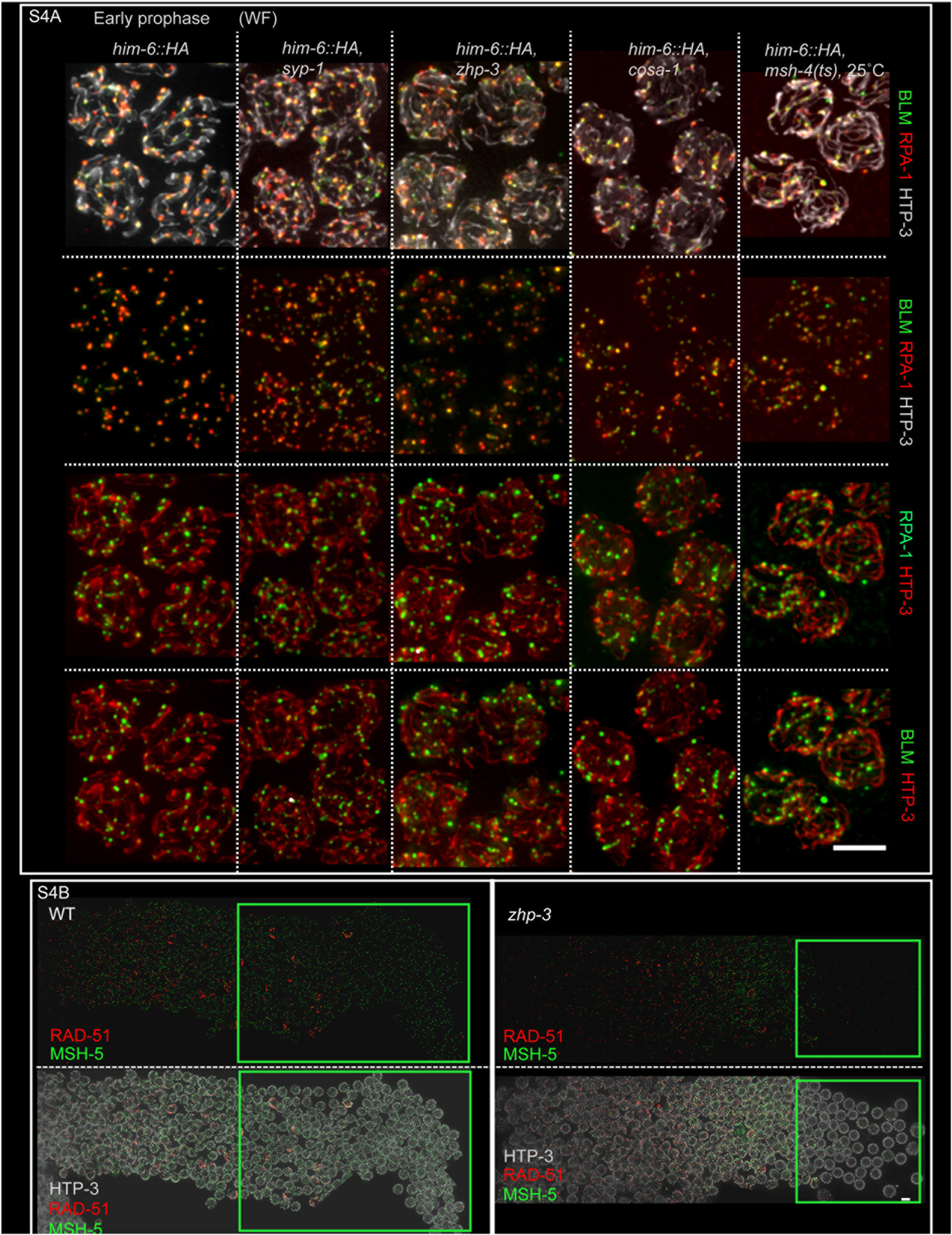
(A) BLM and RPA-1 colocalize in early prophase in SC-CR- and CO-defective mutants. Representative early prophase nuclei from spread gonads probed for BLM::HA, RPA-1 and HTP-3 from WT, null mutants of *syp-1*, *cosa-1*, and *zhp-3*, and a temperature-sensitive *msh-4* mutant. *msh-4ts; him6::HA* animals were grown at 25°C and analyzed 16 hours post L4 larval stage; all other genotypes were assessed under standard conditions (20°C and 24h post L4). Despite higher numbers of recombination foci than in WT, all BLM foci co-localize with RPA-1 foci during early prophase in all these CO-deficient mutants. Please note changes of colors for different markers in the different panels to visualize co-localization. Scale bars represent 5 μm. **(B) DNA repair proteins disappear from chromosomes at the delayed early-to-late prophase transition in a CO-defective mutant.** Images of WT and *zhp-3* mutant germ lines, probed for RAD-51 (red), MSH-5 (green) and HTP-3 (grey), demonstrating the loss of both MSH-5 and RAD-51 foci in the *zhp-3* mutant upon (delayed) transition to late pachytene. Scale bar represents 5 μm.

**Fig S5 (related to Figure 5).**
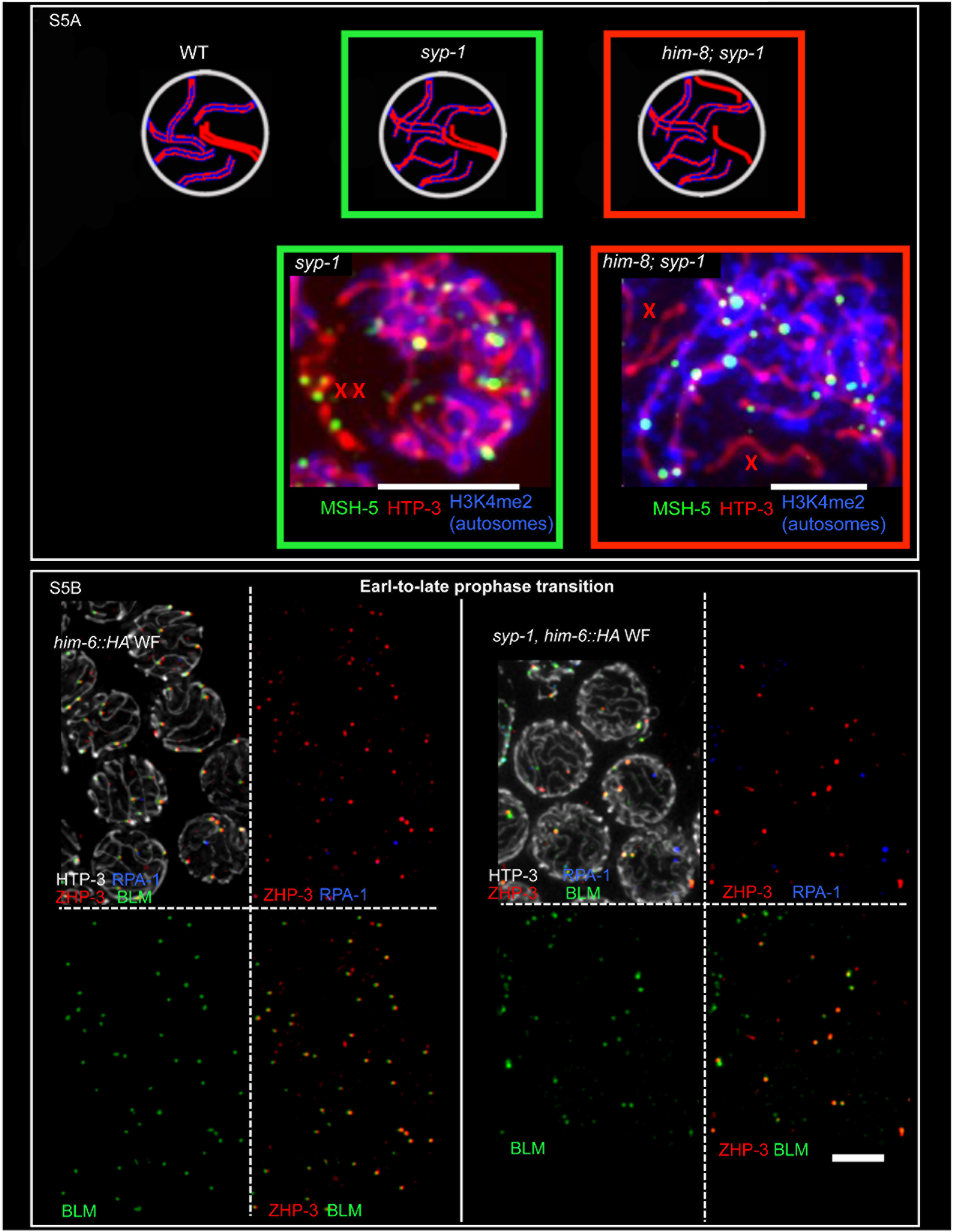
(A) Detection of MSH-5 foci at DSB repair sites in the absence of the SC-CR requires homolog engagement. Top: Schematic of chromosome arrangements in early prophase in WT, (where all homologous chromosomes are paired and synapsed), in the *syp-1* mutant (in which homologs engage in strong pairwise associations in the vicinity of pairing centers (PC), but alignment is not stabilized by synapsis) and in the *him-8; syp-1* double mutant (in which the autosomes engage in unstable PC-mediated pairing, but the X chromosomes never pair). The autosomes are indicated in blue and red, the X chromosomes in red alone. HIM-8 is required for X chromosome pairing and synapsis. Bottom: representative images of early prophase spread nuclei from *syp-1* (top) and *him-8; syp-1* mutant gonads, probed for MSH-5, HTP-3, and di-methylated Lysine 4 on Histone 3 (H3K4me2), which specifically marks the autosomes, but not the X chromosomes. Multiple MSH-5 foci are detected on the X chromosomes in the *syp-1* single mutant, where the X chromosome are able to pair, but MSH-5 foci are absent from the unpaired X chromosomes in the *him-8; syp-1* double mutant. Scale bars represent 5 μm. **(B) ZHP-3 localizes to recombination sites lacking RPA in the SC-CR mutant.** Fields of nuclei from the early-to-late prophase transition regions of WT and *syp-1* spread gonads, probed for HTP-3 (red), BLM::HA (green), RPA-1 (blue) and ZHP-3, a conserved E3 ligase essential for crossing-over (red). Using spreading procedures, SC-associated ZHP-3 that is normally detected in whole-mount preparations of WT gonads is largely washed out, while recombination-site-associated ZHP-3 remains at sites lacking RPA. ZHP-3 is also detected at DNA repair sites in *syp-1* mutant gonads from the early-to-late-prophase transition onward, also at sites lacking RPA. Scale bar represents 5 μm.

